# Proteomic analysis of *Mycobacterium tuberculosis* lacking a putative short-chain dehydrogenase (Rv0148)

**DOI:** 10.1101/2025.08.20.671283

**Authors:** Gunapati Bhargavi, Anbarasu Deenadayalan, Kannan Palaniyandi, Selvakumar Subbian

**Affiliations:** Department of Immunology, Indian Council of Medical Research-National Institute for Research in Tuberculosis, Chetpet, Chennai, India; Public Health Research Institute, New Jersey Medical School, Rutgers University, Newark, NJ, USA

**Keywords:** mycobacterium, ESX secretion system, peptidoglycan, stress response, STRING analysis, protein-protein interactions, LC/MS analysis

## Abstract

*Mycobacterium tuberculosis* (*Mtb*) is an intracellular pathogen that survives in host cells by resisting hostile antimicrobial defenses. However, the molecules and mechanisms that contribute to *Mtb*’s intracellular survival are not fully understood. We have previously reported that Rv0148, a putative short-chain dehydrogenase/reductase, plays a significant role in *Mtb* stress response and virulence in in vitro and in vivo models. To further understand the role of Rv0148 in regulating global functions of *Mtb*, we performed comparative proteomic analysis between pathogenic wild-type (WT) and *Δrv0148* mutant strains. Our mass spectrometry-based proteomics approach identified a total of 738 and 469 proteins, respectively, in the WT and *Δrv0148* mutant, with distinct expression patterns. Gene Ontology analysis revealed significant enrichment of proteins involved in biological processes such as resistant to host immune response and protein homeostasis in *Δrv0148* mutant, while peptidoglycan biosynthesis and ribosomal metabolism pathways were downregulated. Further network analysis revealed dysregulation of proteins involved in bacterial stress response, cell wall components, ribosomal and secretory proteins, suggesting impaired translational machinery in *Δrv0148* mutant. Functional categorization of differentially regulated proteins in *Δrv0148* mutant showed broad reprogramming in intermediary metabolism, stress adaptation, and secretion. These findings indicate that Rv0148 functions as a global regulatory node, which influences remodeling of cell wall components and bacterial physiology, potentially balancing survival and stress adaptation mechanisms in *Mtb*.

**IMPORTANCE:** *Mycobacterium tuberculosis* (*Mtb*), the causative agent of tuberculosis (TB), is a notorious pathogen that can resist the hostile host environment to survive intracellularly and to cause disease. However, the molecular determinants that contribute to *Mtb*’s adaptation to resist the host-imposed stress conditions are not fully understood. Previous in vitro and in vivo studies have shown that Rv0148, a putative short-chain dehydrogenase/reductase, is involved in *Mtb* stress response and virulence. In this study, the genome wide proteomic profile of *Mtb* mutant lacking Rv0148 (*Δrv0148*) was investigated. Compared to the wild type Mtb strain, striking changes in proteome profile of *Δrv0148* mutant was noted. Proteins involved in the ESX secretion system, stress response, ribosomal protein metabolism and cell wall components were significantly affected in the *Δrv0148* mutant. The impact of these changes in biological functions that link Rv0148’s role in *Mtb*’s adaptation to stress conditions is discussed.

## INTRODUCTION

Tuberculosis (TB), caused by *Mycobacterium tuberculosis* (*Mtb*), is the top killer among infectious diseases of humans worldwide. The World Health Organization (WHO) has estimated 10.8 million new TB cases and 1.09 million deaths globally in 2024 (1). The pathogenesis of TB is driven by a highly complex and intricate host-pathogen interactions at the site of infection and systemically, leading to a spectrum of clinical outcomes ranging from asymptomatic latent infection (LTBI) to full-blown symptomatic TB disease (2, 3). Soon after entering into the lungs, *Mtb* is engulfed by the host antigen presenting cells (APC), such as macrophages that attempt to destroy the pathogen through various mechanisms, including the production of reactive oxygen and nitrogen species (ROS and RNS) (4–6). However, *Mtb* can evade these antimicrobial responses of APCs and replicate intracellularly by regulating its metabolic and structural adaptation (7, 8). For example, studies on the *DosR* regulon, which includes genes regulated by DosR protein in response to hypoxia and exposure to ROS and RNS, has provided valuable insights into how *Mtb* survives in the oxygen-limited and stressful environments of granulomas, which contributes to better understanding of TB pathogenesis (9–12). In addition to the canonical stress-responsive proteins such as DosR proteins, there are several “putative” *Mtb* proteins such as the alkylhydroperoxidases (Ahp), which catalyze oxidation-reduction and detoxification of peroxides, are crucial for regulating bacterial cellular metabolism and maintain redox balance, which would enable *Mtb* to survive amid various stress and antimicrobial activities of the host immune system (13, 14). For example, *Mtb* uses oxidoreductases to neutralize ROS produced by activated macrophages and contributes to maintaining the redox balance essential for the intracellular metabolism, survival and replication of *Mtb* (15–17). However, studies on the characterization of *Mtb* Ahp genes are very limited, although knowledge on the functional aspects of these molecules would help in understanding how *Mtb* adapts, survives and establishes infection inside the host. Previously, we reported that *rv0148* encodes a multifunctional short-chain dehydrogenase/reductase (SDR) that plays an important role in *Mtb* survival and persistence in the host through its effect on bacterial stress management, and immune evasion (17, 18). We showed that compared to the WT, the *Δrv0148* strain was significantly more susceptible to killing by various stress-inducing agents in vitro, including peroxidase, ROS, RNS and metal ions. However, the *Δrv0148* strain displayed a higher tolerance to several first-line and second-line anti-TB drugs, compared to the WT strain. Furthermore, the mutant strain was attenuated for virulence and showed reduced intracellular survival in THP-1 macrophages, and in the lungs and spleen of infected guinea pigs (17, 18). In addition, macrophages infected with *Δrv0148* strain expressed elevated levels of pro-inflammatory cytokines, such as TNF, compared to WT, suggesting that Rv0148 may play a role in modulating the host immune responses (17–19). However, studies on the genome wide characterization of proteins that were differentially expressed in *Δrv0148*, compared to WT strain are currently lacking.

Proteomics, a comprehensive genome-wide analysis of proteins, has emerged as a powerful approach to delineate and understand the molecular mechanisms underlying TB pathogenesis (20–24). By enabling the identification and quantification of numerous proteins, and interaction networks, proteomics provide intricate details about host-*Mtb* interactions and bacterial adaptation strategies to survive, replicate and persist in the hostile host environment, (20–24).

Building on our previous findings, we applied a proteomics-based bioinformatic analysis in this study to further investigate the role of Rv0148 in TB pathogenesis. Our data show significant alterations in bacterial pathways related to ribosomal function, the secretory system, stress response, protein acetylation, and peptidoglycan biosynthesis in *Δrv0148*, compared to the WT strain. We observed modulation of protein-protein interaction link between acetylated proteins and components of the ESX secretory system in the *Δrv0148* mutant. The marked upregulation of ESX proteins in *Δrv0148* suggests that type VII secretion system contribute to the altered phenotype of this mutant in vitro and in vivo. Overall, these findings underscore the multifaceted role of Rv0148, particularly in regulating *Mtb* stress response, protein secretion, and cellular homeostasis.

## MATERIALS AND METHODS

### Bacterial Strains, Culture Conditions and Chemicals

The *Mtb* H37Rv (WT) and *Δrv0148* strains were grown to mid-log phase (OD600 ∼ 0.8) at 37 ^ο^C in Middlebrook 7H9 broth (BD Difco^TM^, Franklin Lakes, NJ, USA) supplemented with 0.2% glycerol, 0.05% polysorbate 80, and OADC (oleic acid, bovine albumin, dextrose, and catalase) (BD Difco^TM^, Franklin Lakes, NJ, USA). The *Δrv0148* strain was created and validated in our previous studies as described (17). Hygromycin (50 μg/ml) was added in the broth to grow *Δrv0148* strain. All chemicals and reagents used in this study were purchased from Millipore-Sigma (Sigma-Aldrich, Inc, St. Louis, MO, USA) until specified otherwise.

### Protein Extraction and Precipitation

Cultures of WT and *Δrv0148* strains were centrifuged at 4,000 rpm for 10 minutes at 4 °C, and the bacterial pellets were washed once with ice-cold phosphate-buffered saline (PBS) to remove residual media components and resuspended in ice-cold lysis buffer containing 50 mM Tris (pH 8), 20% glycerol, 300 mM NaCl, 10 mM imidazole, 0.5 mM phenyl methyl sulphonyl fluoride (PMSF) and a protease inhibitor cocktail. The bacterial suspension was kept on ice and sonicated using an ultrasonic homogenizer for 28-30 cycles, with each cycle consisting of a 1-minute pulse followed by a 1-minute interval. The resultant lysate was centrifuged at 10,000 rpm for 30 minutes at 4 °C, to remove the cell debris and unbroken cells. The supernatant containing whole cell lysate was transferred to a fresh tube and subjected to protein precipitation using trichloroacetic acid (TCA) as follows: In a 1.5 mL microcentrifuge tube, 1 mL of the whole cell lysate was mixed with 250 µL of 100% TCA and vortexed vigorously for 1 minute to ensure thorough mixing. The sample was incubated overnight at 4 °C to allow protein precipitation. The samples were centrifuged at 14,000 rpm for 10 minutes at 4 °C, and the supernatant was carefully discarded. The pellet was washed twice with 250 µL of cold acetone to remove residual TCA and other contaminants. Finally, the protein pellet was air-dried at room temperature for 5 minutes and reconstituted in sterile distilled water. The protein concentration was determined by using the bicinchoninic acid (BCA) assay kit following the manufacturer’s protocol (Thermo Fisher Scientific, CA, USA). The integrity and yield of extracted protein were confirmed by SDS-PAGE prior to downstream processing.

### SDS-PAGE and Gel Processing

To determine the integrity and quality, 20μg of extracted proteins per sample was resolved on 12.5% SDS-PAGE gels. The gels were incubated for 3 hours with a staining solution containing 0.25% Coomassie blue R-250, 40% methanol, 10% glacial acetic acid, and 50% distilled water. The stained gels were destained using a solution composed of 45% methanol, 10% glacial acetic acid, and 45% water. The destaining solution was changed periodically until the Coomassie blue was completely removed from the gels. Finally, the gels were rinsed with water and protein bands of interest were excised carefully using a sterile scalpel. The excised gel pieces were diced into ∼1 mm^3^ cubes, transferred into 1.5 mL microcentrifuge tubes and washed with Milli-Q water to remove residual stains and contaminants. Briefly, the gel fragments were vortexed at 1,500 rpm for 1 hour, followed by removal of the wash solution and a repeat wash under identical conditions. After washing, the gel pieces were spun briefly at 3,000 rpm for 20 seconds, and the supernatant was carefully discarded. Subsequently, 300 µL of a 1:1 (v/v) mixture of 100 mM ammonium bicarbonate and acetonitrile was added to each tube, followed by vortexing at 1,500 rpm for 30 minutes to further destain the gel pieces. This was followed by the addition of 500 µL of 100% acetonitrile and incubation at room temperature with gentle vortexing until the gel pieces turned transparent. The acetonitrile was completely removed, and tubes were left open for 20 minutes to allow residual solvent to evaporate. and the gel pieces were stored at -20 ^ο^C.

### In-Gel Trypsin Digestion and Peptide Extraction

For enzymatic digestion, the dried gel pieces were rehydrated with 50 µL of sequencing-grade trypsin solution (13 ng/µL in 50 mM ammonium bicarbonate) and incubated on ice for 30 minutes. The buffer volume was checked and, if needed, additional trypsin buffer was added to ensure complete rehydration of the gel pieces. After absorption of the enzyme solution into the gel matrix, the samples were incubated overnight at 37°C with gentle shaking at 300 rpm to facilitate protein digestion. Following digestion, the trypsin-containing supernatant was carefully transferred to microcentrifuge tubes. To extract remaining peptides from the gel, an extraction buffer (volume approximately twice that of the trypsin buffer, typically 100 µL) was added to the gel pieces, followed by vigorous vortexing for 20 minutes. The resulting supernatant was pooled with the previously collected trypsin digest. The combined peptide extracts were dried completely using a SpeedVac concentrator and stored at −20°C. Prior to LC-MS analysis, dried peptide samples were reconstituted in 30 µL of 3% acetonitrile.

### Liquid Chromatography–Mass Spectrometry (LC-MS)

The peptides were analyzed using a nano ACQUITY UPLC chromatographic system (Waters, Manchester, UK) equipped with MassLynx4.1 SCN781 software for data acquisition. Chromatographic separation was performed using a Symmetry® C18 trap column (180 μm × 20 mm, 5 μm) and an HSS T3 C18 analytical column (75 μm × 200 mm, 1.8 μm). (Waters, Manchester, UK). The separation was carried out in reverse-phase mode, with a mobile phase consisting of solvent A (0.1% formic acid in water) and solvent B (0.1% formic acid in acetonitrile). The gradient program was as follows: starting conditions were 99% solvent A and 1% solvent B, held for 3 minutes. The composition was then linearly changed to 60% solvent A and 40% solvent B by 43 minutes, and further to 20% solvent A and 80% solvent B by 46 minutes, which was maintained until 50 minutes. The gradient was then returned to the initial conditions of 99% solvent A and 1% solvent B for 51 minutes and held for 60 minutes. The flow rate was set at 300 nL/min, with the column temperature maintained at 35 °C and the auto sampler temperature at 4 °C. The peptides eluted from the chromatography column were analyzed through mass spectrometry (MS).

### Mass Spectrometry Acquisition and Protein Identification

All the MS runs were performed using ion mobility-enabled separation on a Synapt G2 High-Definition MS™ System (HDMSE System; Waters, Manchester, UK). Data acquisition was conducted positively, with a nano electrospray ionization (nano ESI) capillary voltage set at 3.4 kV. The sample cone voltage was 40 V, and the extraction cone voltage was 4 V. The ion mobility spectrometry (IMS) gas (N2) flow was maintained at 90 mL/min. The IMS T-Wave™ pulse height was set at 40 V, and the IMS T-Wave™ velocity was 800 m/s, with IMS ramping between 8 V and 20 V. The system operated in resolution mode, and data was acquired in a continuum format. The collision energy was ramped from 20 eV to 45 eV. Raw MS data was processed using Progenesis QI for Proteomics V4.2 (Non-Linear Dynamics, Waters) as described previously (25, 26). Protein identification was performed using *Mtb* databases sourced from UniProt (https://www.uniprot.org/) reviewed entries (retrieved on 31.11.2021). The identification criteria included ≥1 fragment per peptide, ≥3 fragments per protein, and ≥1 peptide per protein. Cysteine carbamidomethylation was considered a variable modification, while methionine oxidation was treated as a fixed modification. For each of the identified and annotated Mtb protein, the confidence score was determined with a cutoff false discovery rate (FDR) of 4%, with allowance for 1 missed cleavage.

### Proteomics Data Analysis

Based on the MS data on identified Mtb proteins with significant confidence scores, separate protein lists for WT H37Rv and *Δrv0148* strains were generated. The raw expression data of proteins in WT and *Δrv0148* strains is presented in Supplementary Table-1. These lists were further annotated to assign Mtb protein IDs (Rv numbers) and analyzed using Venny 2.1.0 to identify unique- and commonly-expressed proteins between WT and *Δrv0148* strains. To determine differentially expressed proteins, expression ratios were calculated by dividing the confidence scores of WT and *Δrv0148* strains. Proteins with expression ratios >2 were classified as upregulated, and those with ratios <0.5 were considered downregulated in *Δrv0148* compared to the WT strain. Functional classification of these proteins was performed using ShinyGO (v0.75), a software that determines the significantly regulated biological pathways and gene ontology (GO) terms by analyzing enriched versus background protein sets and calculating FDR based on fold change and p-values, which indicate the statistical significance of enrichment for each functional category. Further, STRING 12.0 was employed to cluster the upregulated and downregulated proteins based on predicted functional protein-protein interactions (PPI) and categorize them based on functional roles. Finally, GraphPad Prism 10 was used to visualize the expression patterns, highlighting proteins involved in pathways that were distinctly upregulated and/or downregulated in WT and/or *Δrv0148* strains.

## RESULTS

### Comparative Proteomic Analysis of WT H37Rv and *Δrv0148* Strains

To assess the impact of *rv0148* deletion on genome wide expression of *Mtb* proteome, we performed a comparative proteomic analysis between the WT and *Δrv0148* mutant strains. As shown in Fig. 1A and Supplementary Table-2, among a total of 887 proteins that were differentially expressed across both strains, 149 were expressed only in *Δrv0148*, compared to 418 unique proteins in the WT, while 320 proteins were commonly expressed in both strains (Fig.1A and B). Distinct expression patterns were observed among the proteins expressed both in the WT and *Δrv0148* strains, marked by divergent confidence scores, as shown in the Volcano Plot in Fig.2A. Compared to the WT, expression of 10 proteins (Rv2226, Rv2614c, Rv0282, Rv2703, Rv2477c, Rv2572c, Rv0120c, Rv0480c, Rv2971 and Rv1872c) were upregulated, while 13 proteins (Rv2539c, Rv0896, Rv0009, Rv2461c, Rv1475c, Rv0643c, Rv2839c, Rv1094, Rv2750, Rv0079, Rv2175c, Rv1390 and Rv0292) were downregulated by more than 10-folds in *Δrv0148* strain (Fig.2B). The nature and putative function of these differentially regulated proteins indicates that *rv0148* may act as a regulatory node in Mtb, influencing multiple cellular pathways, potentially through redox-sensitive transcriptional or post-translational mechanisms.

**Fig. 1.**
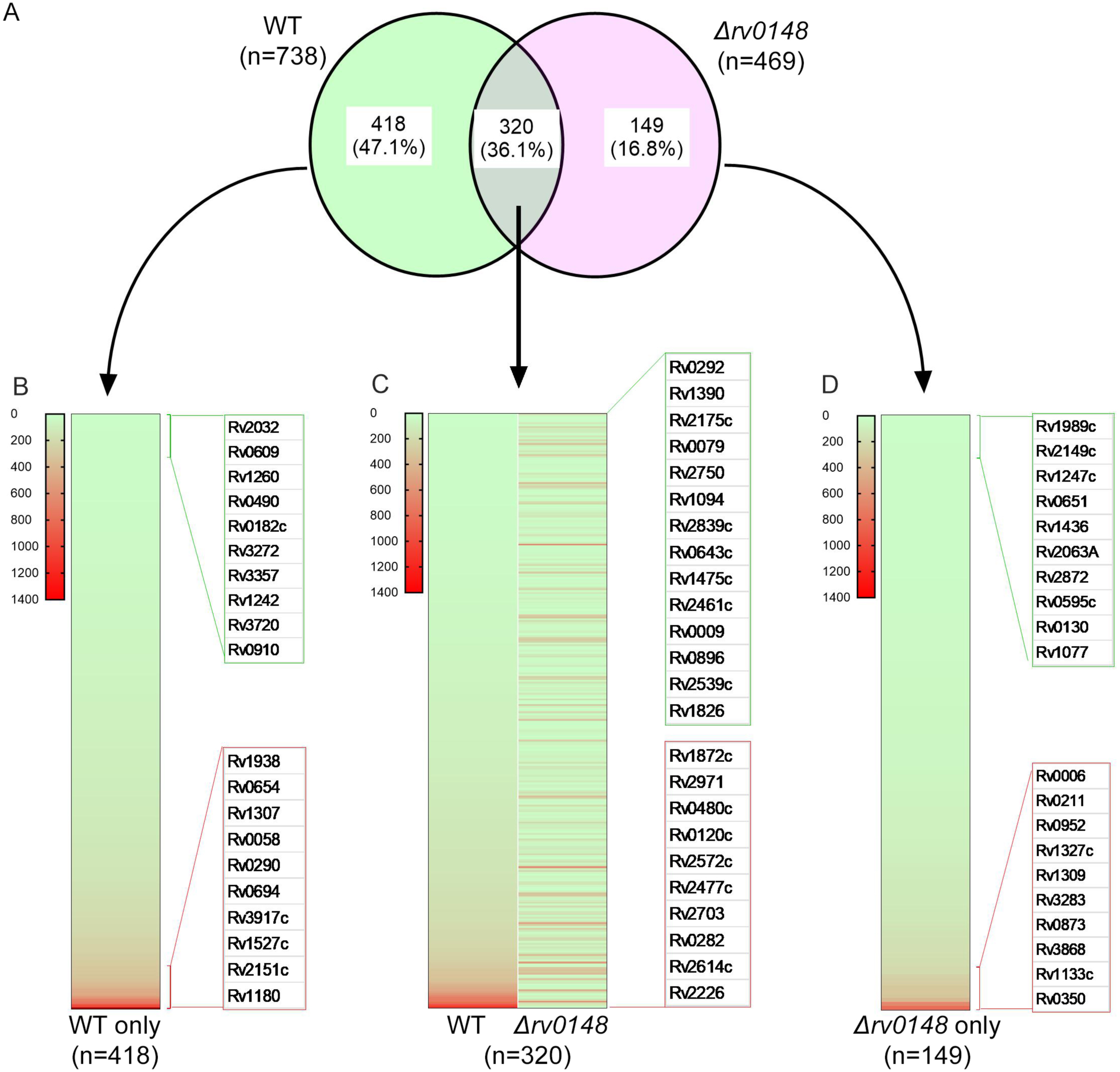
Comparative proteomic analysis of wildtype (WT) and *Δrv0148* mutant strains. (**A**) Venn diagram showing the number of unique and shared proteins between WT and *Δrv0148* mutant. A total of 887 proteins were detected, which includes 418 unique to WT, 149 unique to *Δrv0148*, and 320 common to both strains. (**B**) The confidence score-based heatmap showing the expression profile of 418 proteins uniquely regulated in WT. The top 10 up- and down-regulated proteins are listed in red and green boxes, respectively. (**C**) Heatmap showing the expression profile of 320 proteins commonly regulated in WT and *Δrv0148* mutant. Proteins that are up- or down-regulated in *Δrv0148* mutant, compared to WT are listed in red and green boxes, respectively. (**D**) Heatmap showing the expression profile of 149 proteins uniquely regulated in *Δrv0148* mutant. The top 10 up- and down-regulated proteins are listed in red and green boxes, respectively. Scale bar in B-D corresponds to confidence scores ranging from 1400 (highly upregulated) to 0 (highly downregulated). The heatmap was created using GraphPad Prism 10.

**Fig. 2.**
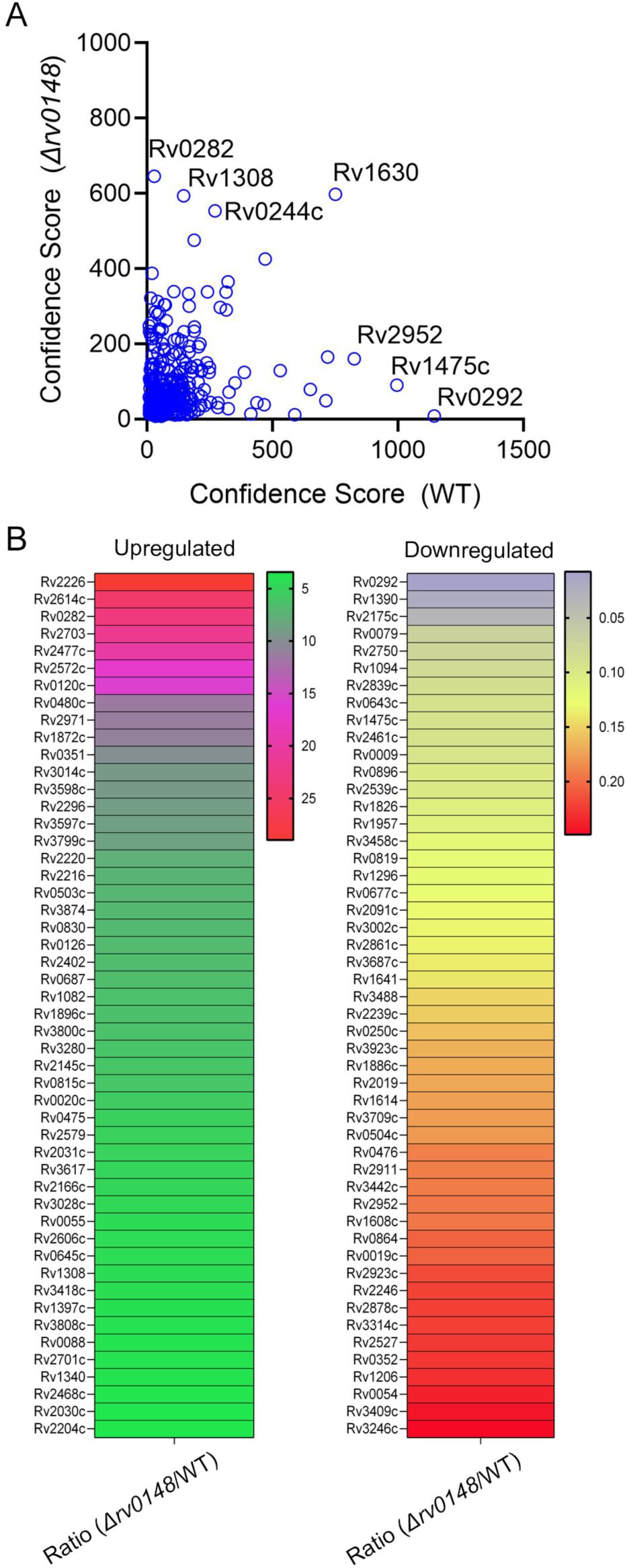
Differential expression profile of proteins in the WT and *Δrv0148* mutant strains. (**A**). Volcano plot showing the expression patten of commonly regulated proteins between WT versus *Δrv0148* mutant. Each point represents the confidence score of proteins in the *Δrv0148* mutant (y-axis) compared to the WT (x-axis). The plot illustrates the overall distribution and variation of proteins based on confidence scores, highlighting proteins that are more upregulated in the WT or *Δrv0148* mutant. (**B**). Heatmap of top 50 upregulated proteins in *Δrv0148* mutant, relative to WT. (**C**). Heatmap of top 50 downregulated proteins in *Δrv0148* mutant, relative to WT. The ratio of protein expression levels in *Δrv0148* mutant, compared to WT was used to build the heat map. Color intensity reflects the fold change in expression, with the gradient scale indicating increasing expression from green/ pink/ red (**B**) or blue/ yellow/ red (**C**). The volcano plot and heatmaps were created using GraphPad Prism 10.

### GO analysis of Upregulated Proteins in the Δ0148 mutant

To further understand the functional implications of the proteins upregulated in the *Δrv0148,* compared to the WT, we performed gene ontology (GO) analysis using ShinyGO. The significantly enriched GO terms (FDR<0.05) were categorized into three main domains: (i). biological processes, (ii). cellular compartments, and (iii). molecular functions (Fig.3). In the biological processes category, a total of 50 significantly enriched GO was identified (Fig.3A). Prominent among these were those associated with stress response and virulence, such as host immune response, pathogenesis, cellular response to starvation, and response to chemical or cellular stimuli. These enrichment findings suggest that deletion of Rv0148 impacts *Mtb* adaptive mechanisms to survive under hostile stress conditions in the host. Additional GO enriched terms, such as protein folding, cell communication, cellular nitrogen compound metabolic process, and regulation of primary metabolic process, indicate a broad shift in reprogramming of cellular functions toward managing protein homeostasis in *Δrv0148* strain (Fig.3A).

**Fig. 3.**
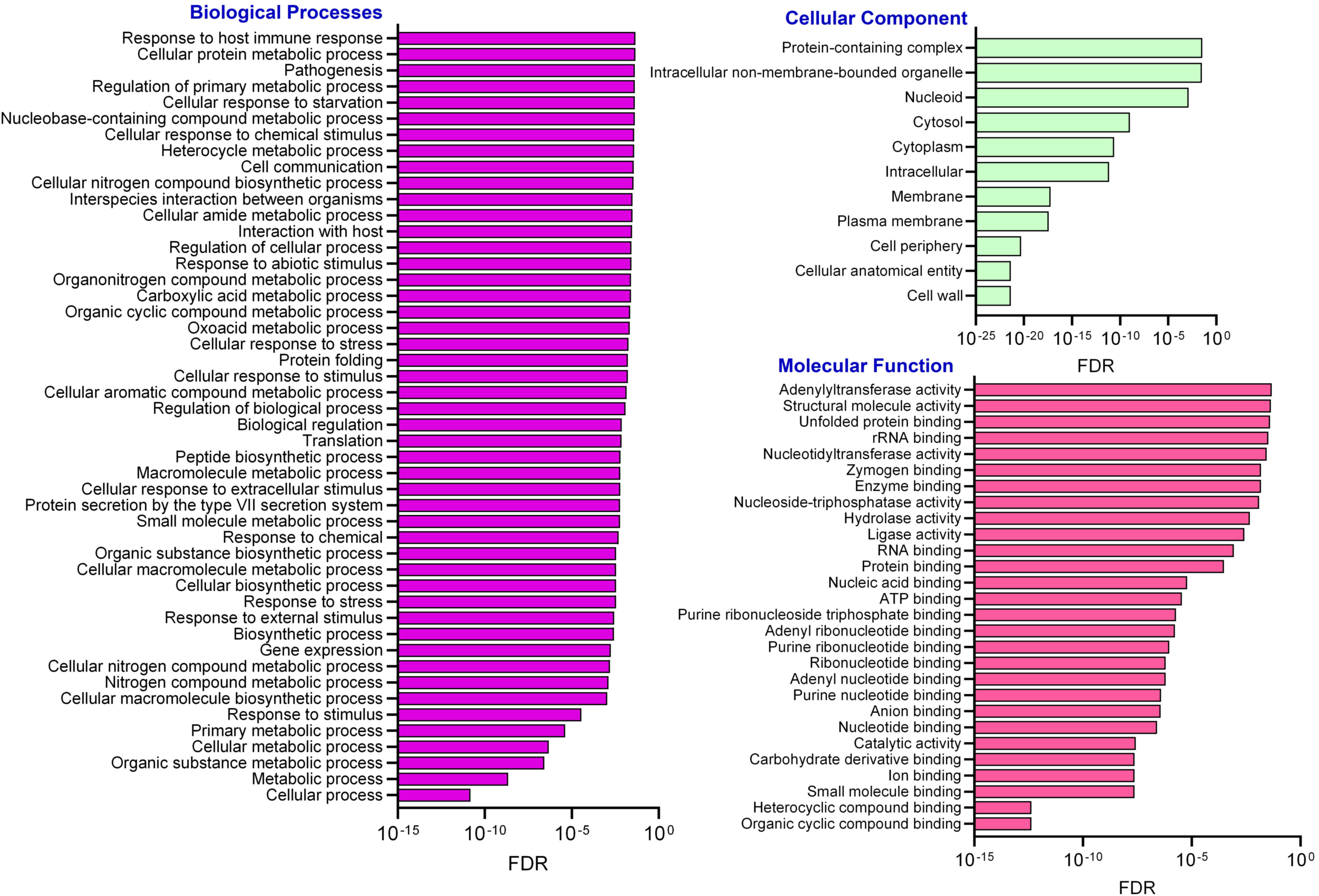
GO enrichment analysis of significantly upregulated proteins in *Δrv0148* mutant compared to WT. The bar plots show enriched GO categories in biological processes, cellular component and molecular functions. Enrichment analysis was performed using upregulated proteins in *Δrv0148* mutant relative to WT. The x-axis represents the FDR, plotted on a log scale. Lower FDR values indicate higher statistical significance of enrichment. The heatmaps were created using GraphPad Prism 10.

Analysis of the cellular component category revealed that the upregulated proteins in *Δrv0148* strain were predominantly localized to protein-containing complexes, nucleoid, cytosol, and intracellular non-membrane-bounded organelles (Fig.3B). This enrichment pattern aligns with increased transcriptional and translational activity, as well as potential structural reorganization, reflecting a compensatory response in *Δrv0148* strain to maintain cellular integrity under stress. In the molecular function category, analysis revealed that the upregulated proteins were enriched in binding and catalytic enzymatic activities (Fig.3C). Functional categories such as adenylyl transferase activity, unfolded protein binding, rRNA binding, ATP binding, and nucleoside-triphosphatase activity were significantly represented. These results suggest upregulation of processes involved in nucleotide metabolism, protein folding, and ribonucleotide processing, likely supporting cellular adaptation of *Δrv0148* mutant. Importantly, a remarkable interaction among various pathways and networks comprised of upregulated protein was noted, which indicate coordinated regulation different functions perturbed in the *Δrv0148* mutant (Supplementary figure S1A).

### GO analysis of Downregulated Proteins in the Δ0148 mutant

Interestingly, a wide range of biological functions were downregulated in *Δrv0148* compared to the WT, particularly those related to biosynthesis and regulation pathways (Fig.4A). Highly enriched GO terms in the biological processes of *Δrv0148* strain include the regulation of molecular function, carboxylic acid metabolic process, cellular response to stress, and negative regulation of macromolecule biosynthetic process. In addition, cellular processes linked to translation, protein metabolic process, gene expression, homeostasis, and nitrogen compound biosynthesis were also significantly suppressed, indicating a broad downregulation of stress response and core metabolic systems in *Δrv0148,* compared to the WT *Mtb*.

**Fig. 4.**
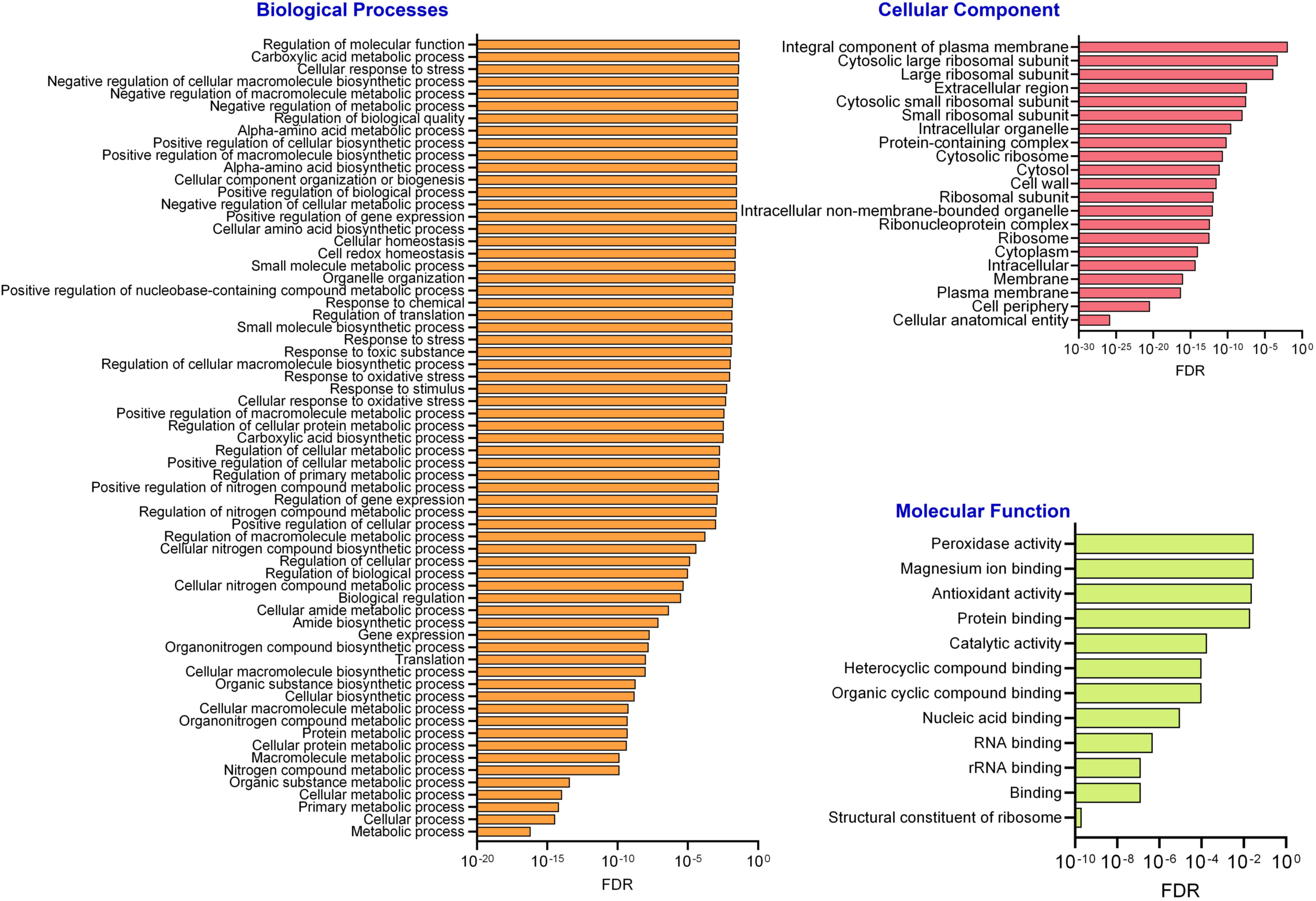
GO enrichment analysis of significantly downregulated proteins in *Δrv0148* mutant compared to WT. The bar plots show enriched GO categories in biological processes, cellular component and molecular functions. Enrichment analysis was performed using downregulated proteins in *Δrv0148* mutant relative to WT. The x-axis represents the FDR, plotted on a log scale. Lower FDR values indicate higher statistical significance of enrichment. The heatmaps were created using GraphPad Prism 10.

Most of the downregulated proteins enriched in the cellular component category of *Δrv0148* strain were primarily associated with ribosomal subunits, including large and small ribosomal subunits, cytosolic ribosomes, protein-containing complexes, cytosol, and intracellular organelles (Fig.4B). These findings suggest a significant suppression of ribosomal biogenesis in *Δrv0148* affecting translational activity. Additional components, such as the extracellular region, the plasma membrane, and the cell periphery, were also enriched in *Δrv0148* strain, hinting at the broader effects on bacterial cell structure, potentially impacting the secretion pathways. Furthermore, the most significantly suppressed molecular functions in *Δrv0148* strain include peroxidase activity, magnesium ion binding, antioxidant activity, protein binding, and catalytic activity (Fig.4C). Several biomolecule-binding activities, including nucleic acid binding (e.g., RNA binding) and heterocyclic compound binding, were also reduced in *Δrv0148* strain. These patterns imply a decrease in enzymatic detoxification, oxidative stress defense, and nucleic acid-associated binding functions, pointing to a weakened oxidative stress response in *Δrv0148*, compared to WT. In addition, significant interaction among various pathways and networks involving downregulated protein was noted, which indicate coordinated regulation different functions perturbed in the *Δrv0148* mutant (Supplementary figure S1B).

### Functional Categorization of Differentially Expressed Proteins in *Δrv0148* mutant

To further contextualize the biological significance of differentially regulated proteins in *Δrv0148*, we performed a GO-based functional categorization analysis (Fig.5). Among all differentially regulated proteins in *Δrv0148*, the highest number of upregulated (n=123) and downregulated (n=148) belongs to the cellular anatomical entities category. Consistent with this, 70 of the upregulated and 80 of the downregulated proteins were localized in the cytoplasm, while 81 upregulated and 93 downregulated proteins were localized to the intracellular regions. These observations suggest notable changes in *Mtb* structural protein localization in *Δrv0148*, compared to WT.

**Fig. 5.**
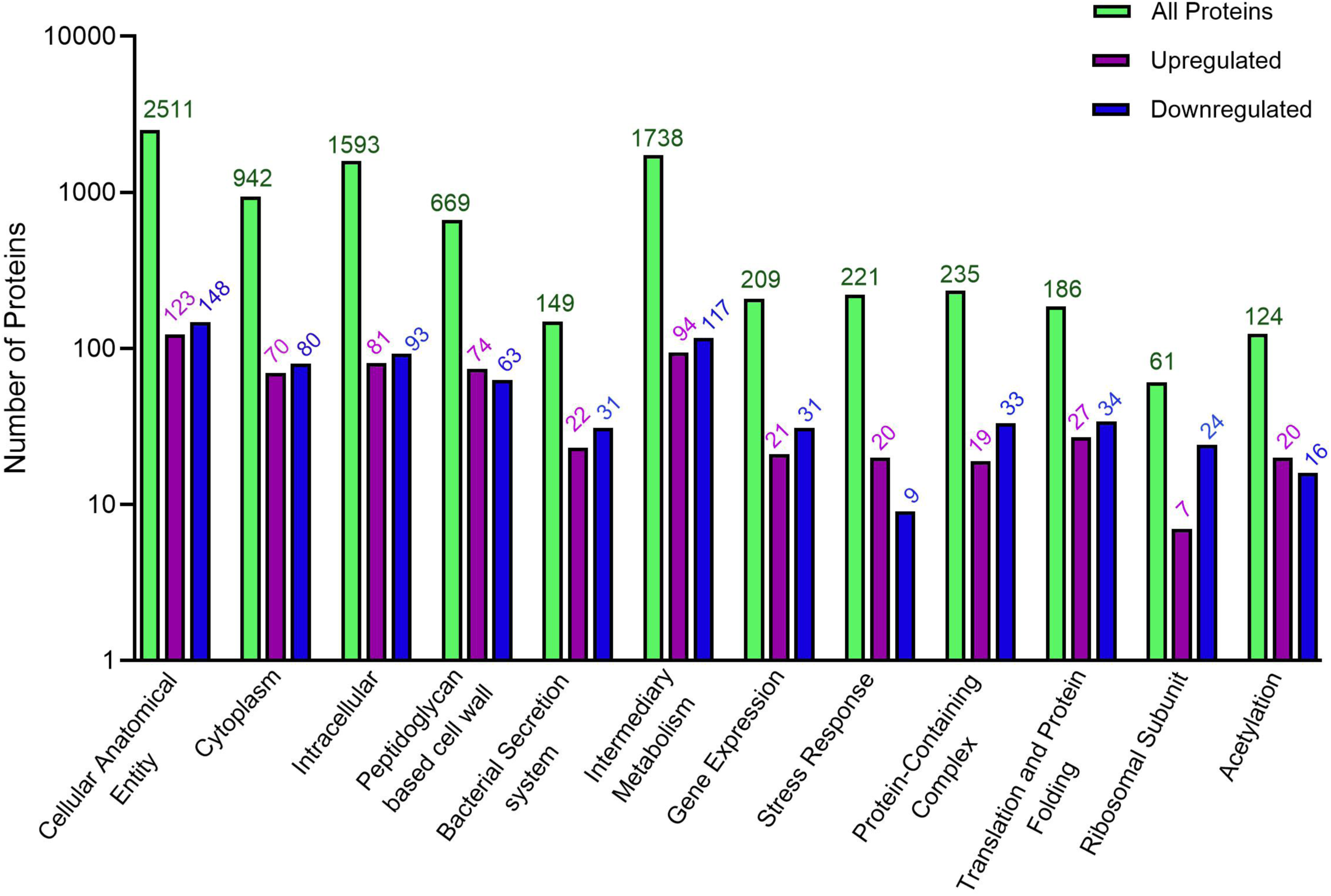
Functional classification of differentially expressed proteins in *Δrv0148* mutant. The bar chart displays the distribution of proteins that are identified across various functional categories in *Δrv0148* mutant, compared to expression levels in WT. The total number of all proteins in each functional category is shown in green (all proteins). Distribution of differentially up and down regulated proteins in *Δrv0148* mutant are shown in purple and blue colors, respectively. Data analysis for functional classification was defined and assigned based on Mycobrowser Mtb H_37_Rv annotation and STRING 12.0 database. The bar graph was created using GraphPad Prism 10.

A significant number of differentially expressed proteins were associated with intermediary metabolism (94 upregulated and 117 downregulated), indicating broad metabolic remodeling. Particularly, a greater number of proteins associated with peptidoglycan-biosynthesis and cell wall processes were upregulated (n=74), compared to downregulation (n=63), which suggests alterations in cell wall composition. This is consistent with impaired adaptation of *Δrv0148* strain to stress and antibiotic pressure. A similar expression pattern was also observed in stress response proteins (20 upregulated and 19 downregulated) and acetylation-related proteins (20 upregulated and 16 downregulated), suggesting that post-translational modifications may contribute to the observed proteomic shifts in *Δrv0148* strain as an adaptation to stress response through modulation of bacterial structural components (Fig.5). Additionally, proteins involved in mycobacterial secretion system were significantly altered (22 upregulated and 31 downregulated) in *Δrv0148* strain, which is consistent with our STRING analysis suggesting modulation of ESX-1 secretion components. Further, proteins involved in translation and protein folding showed differential expression, with notable upregulation of chaperones such as GroES and GroEL1, which are known to be induced under heat shock and oxidative stress (27, 28). Taken together, these proteomic alterations suggest that deletion of *rv0148* impacts multiple functional pathways in *Mtb*, notably those related to bacterial metabolism, adaptation to stress, secretion of virulence factors, and protein homeostasis.

### STRING analysis of highly enriched networks in *Δrv0148* mutant

To further delineate the expression profile of key proteins that are functionally relevant to the phenotype of Δ*rv0148* mutant, we performed protein-protein interactions (PPI) of differentially expression proteins through STRING analysis (Supplementary Fig-S2). The major functional categories that emerged from this analysis included the secretory system, ribosomal proteins, stress, acetylation and peptidoglycan metabolism. These networks were also highly enriched in our functional categorization analysis as shown in Fig-5.

*a). Mtb Secretory System network:* Proteomic profiling of the *Δrv0148* mutant revealed that 15 of the proteins (thrS, eccA3, sigA, aspS, fusA2, lysS, rpsR1, atpA, rpsT, amiC, rpmA, rho, rpoA, greA, rplC and rpoB) involved in bacterial secretory system, including the ESX-1 system (eccA3), were upregulated by >2-fold; among these, 4 proteins (thrS, SigA, aspS, and fusA2) were upregulated by >10 folds. In addition, 10 proteins in this network (rpsG, rbpA, rplD, rpsQ, rplV, yajC, rpmC, gpsI, infC and rpoZ) were downregulated by >2 folds, with rpoZ downregulated by >10-fold in Δ*rv0148* mutant, compared to the WT (Fig.6A). While the upregulated molecules reflect compensatory regulatory mechanisms or shifts in metabolic priorities in response to the *Δrv0148* deletion, the downregulated molecules suggest a broad downregulation of proteins particularly linked to ribosomal function and cellular secretion (Fig.6A and Supplementary table-1).
*b). Mtb Ribosomal protein metabolism network:* In the *Δrv0148* mutant, we observed a notable reduction in the abundance of several 30S and 50S ribosomal proteins compared to WT *Mtb* (Fig.6B). Most prominently, 30S ribosome-associated proteins of Mtb, such as rpsG, rpsD, rpsL, and rpsQ, and rpsT, as well as 50S ribosome-associated protein, rpmC, displayed > 10-fold reduced expression in *Δrv0148* mutant. In contrast, expression of 4 ribosomal proteins (rpsR1, rpsT, rpmA and rplC) were upregulated by >2-fold in *Δrv0148* mutant, compared to the WT (Fig.6B and Supplementary table-1). These results suggest a selective dysregulation of ribosomal protein expression in the absence of *rv0148*, particularly affecting components of the 30S subunit.
*c). Mtb Stress Response network:* Fifteen proteins, including the chaperone proteins groES and groEL1 and heat-shock protein (hspX) were upregulated by >2 fold, while seven other proteins, including sigF, ahpE and katG that are involved in ROS stress were downregulated by >2 fold in *Δrv0148* mutant, compared to the WT (Fig.6C). The pattern of stress response network protein expression in *Δrv0148* mutant suggests lack of key components protecting against specific stress conditions such as ROS, and activation of other compensatory molecules to still protect the bacteria.
*d). Mtb Acetylation Proteins:* Protein acetylation is a critical post-translational modification that regulates protein activity and stability, with context-dependent roles across diverse cellular processes. With our earlier observations linking acetylation with the secretory pathway and stress response, we further explored the relationship between protein acetylation and the ESX secretory system. Interestingly, we observed several ESX system-associated proteins, including esxA, esxB, espB and espF, as well as proteins associated with stress response, such as grpE, cmaA2, pknD, greA, fabG1, thrS and suhB upregulated by >2 fold in *Δrv0148* mutant, compared to WT (Fig.6D and Supplementary table-1). Conversely, expression of 12 proteins, including Rv0250c, Rv0504c, desA2, rpIV, gcvH, secBL, TB18.6, mmpS3, cysA1, hupB, rpsl and espC, were downregulated by >2 fold in *Δrv0148* mutant (Fig.6D).
*e). Peptidoglycan metabolism:* Peptidoglycan provides the structural integrity of the cell wall and survival of *Mtb*. Based on the functional connection between stress response, secretory system and Mtb cell wall integrity, we analyzed the peptidoglycan metabolism proteins. Of the 58 proteins that were differentially regulated by >2 folds, 19 were upregulated and 39 were downregulated in *Δrv0148* mutant (Fig.6E). Among these, grpE, IldD2, Rv0480c, fusA2 and eccA3 were upregulated, while rpoZ and eccE3 were downregulated by >10 fold in the mutant, compared to the WT. These findings highlight the significant perturbations in the stress response and secretion systems that are also linked to peptidoglycan-associated proteins following *rv0148* deletion, suggesting the complex coordination of cell wall components and adaptation mechanisms during stress in Mtb.

**Fig. 6.**
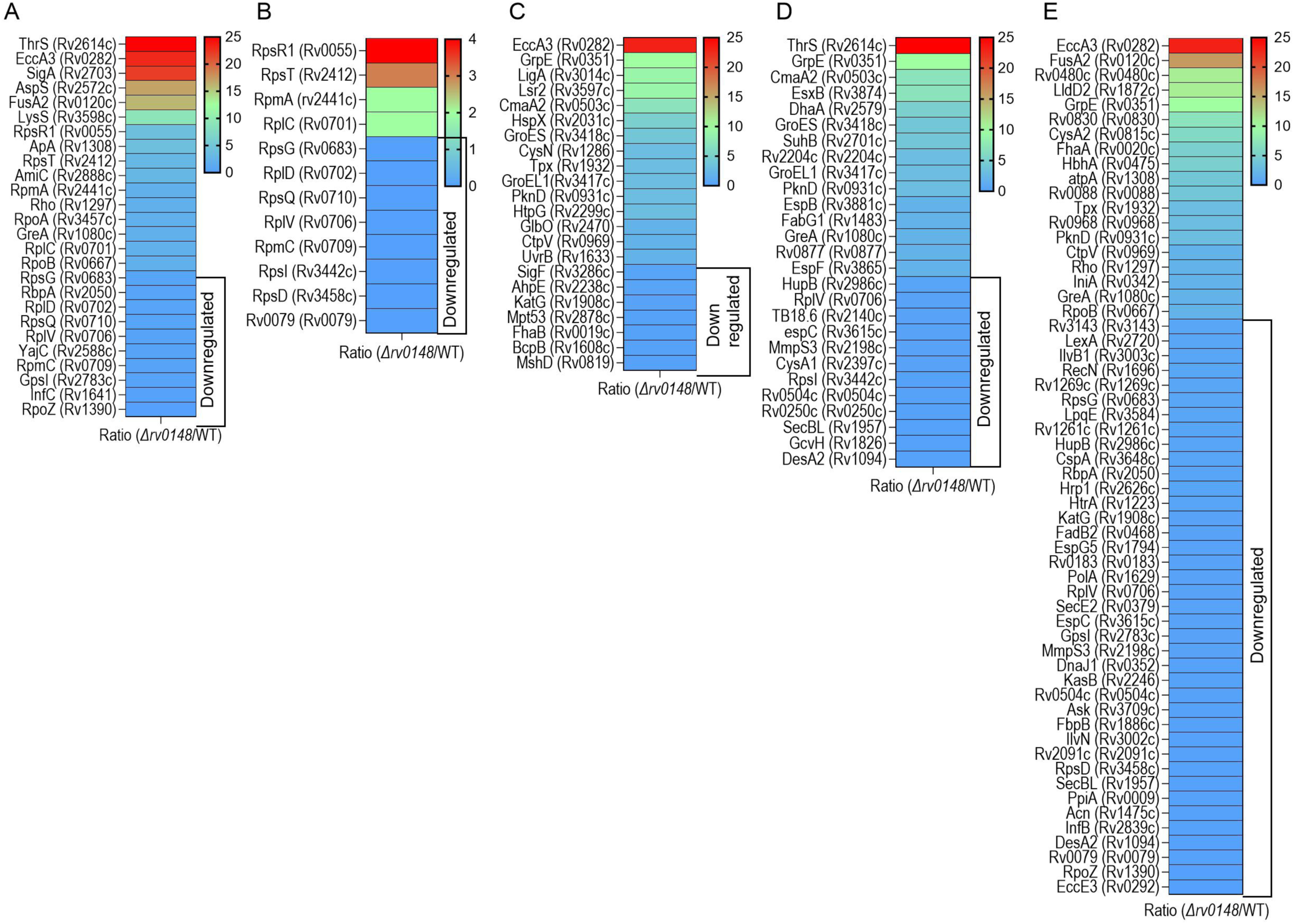
Differential expression of proteins in specific pathways in *Δrv0148* mutant. (**A**). Secretory systems. (**B**). Ribosomal proteins. (**C**). Stress response. (**D**). Acetylation proteins. (**E**). Peptidoglycan metabolism. The ratio of expression levels between *Δrv0148* mutant and WT was used to generate the heat maps in **A**-**E**. Scale bars are represent the ratio of expression. The color intensity represents the highest (red color) or lowest (blue color) expression ratio in the *Δrv0148* mutant. Downregulated proteins are marked with boxes in each of the heat maps. The heatmaps were created using GraphPad Prism 10.

## DISCUSSION

This study provides a comprehensive proteomic dissection of the regulatory role of Rv0148, a putative short-chain dehydrogenase/reductase in *Mtb*. Through high-resolution label-free quantitative proteomics analysis, we demonstrate that deletion of Rv0148 induces a profound reprogramming of the Mtb proteome, affecting over 700 proteins across diverse functional categories, impacting bacterial redox regulation, stress adaptation, and virulence modulation. The proteome profile not only underscores the breadth of Rv0148’s regulatory influence on *Mtb* physiology, but also consistent with its classification within the SDR family, which is known to modulate redox homeostasis and intracellular survival during infection (17–19). We observed that the absence of Rv0148 affects the intracellular redox equilibrium, triggering compensatory activation of stress-responsive pathways and suppression of biosynthetic processes in *Mtb*. This imbalance is reflected in the upregulation of chaperones, such as GroEL1 and GroES, which are pivotal in maintaining proteostasis under oxidative, thermal and chemical-induced stresses (29, 30). For example, GroEL1 has been shown to interact with lipid biosynthesis pathways and contribute to cell wall integrity in *Mtb*, particularly under hypoxic conditions, while GroES prevents aggregation of misfolded proteins and is essential for survival under heat shock and oxidative stress (27, 31). The coordinated upregulation of these chaperones in *Δrv0148* mutant suggests a robust compensatory mechanism to counteract proteotoxic stress. Moreover, the stress-induced expression of other chaperones such as DnaK and ClpB, which assist in refolding and degradation of damaged proteins, further supports the hypothesis that Rv0148 deletion imposes a significant proteostatic burden in *Mtb* (32, 33).

In our analysis, the ESX-1 secretion system, a critical determinant of *Mtb* virulence, was notably modulated in the *Δrv0148* mutant. While some components of ESX-1 system, such as EccA3 were strongly upregulated, others were suppressed, indicating a dysregulated secretion landscape in the mutant. The Mtb ESX-1 facilitates phagosomal escape and cytosolic access via substrates like ESAT-6 and EspB, which disrupt host membranes and modulate innate immune responses (34, 35). Therefore, the unbalanced expression of ESX-1 components in *Δrv0148* suggests that Rv0148 may act upstream of this system, potentially through redox-sensitive transcriptional regulators such as WhiB6, which binds to the promoter of ESX-1 genes and responds to oxidative cues (36, 37). Similarly, EspL, another ESX-1 regulator, modulates WhiB6 expression and is sensitive to redox fluctuations, further implicating Rv0148 in this regulatory axis (38). Additionally, the ESX-1 system is tightly linked to host immune evasion, and its dysregulation may impair Mtb’s ability to modulate macrophage responses, potentially attenuating virulence in the *Δrv0148* strain (39, 40).

We observed ribosomal remodeling as a hallmark of the *Δrv0148* proteome. Particularly, the downregulation of 30S and 50S ribosomal proteins (e.g., RpsG, RpsD, RpsL) aligns with previous reports linking ribosome suppression to oxidative stress and nutrient limitation in *Mtb* (41, 42). This remodeling is a known survival strategy in *Mtb*, allowing the bacterium to conserve energy and prioritize translation of stress-response proteins (41, 43). Interestingly, selective upregulation of ribosomal proteins such as RpsR1 and RplC in *Δrv0148* mutant suggests a nuanced translational reprogramming, rather than global arrest in this strain. This selective modulation may be mediated by ribosome-associated quality control mechanisms, including hibernation-promoting factors (HPFs) and ribosome-splitting enzymes, which have been shown to regulate translation under stress in *Mtb* (44). Moreover, ribosomal protein acetylation and methylation, which influence ribosome stability and mRNA selectivity, may be altered in *Δrv0148* mutant, contributing to the observed proteomic shifts (41).

The proteomics analysis revealed that antioxidant enzymes, including KatG and AhpC, were markedly downregulated in the *Δrv0148* mutant, indicating a compromised detoxification capacity. KatG is a bifunctional catalase-peroxidase, which is essential for neutralizing hydrogen peroxide and activating isoniazid, a frontline anti-TB drug (45, 46). Therefore, downregulation of KatG may render the *Δrv0148* mutant to be more vulnerable to host-derived ROS. This observation is consistent with and supported by our previous studies that showed elevated killing of *Δrv0148* mutant upon exposure to ROS and RNS generating chemicals in vitro (18). Similarly, AhpC, which reduces organic peroxides, also showed reduced expression in *Δrv0148* mutant, further supporting the notion of impaired oxidative stress resistance (47, 48). The downregulation of these enzymes may also reflect a broader failure in redox homeostasis in *Δrv0148* mutant, potentially linked to disrupted NADH/NAD+ ratios or impaired thiol-redox buffering systems. Previous studies have shown that *Mtb* relies on a network of redox-balancing enzymes, including thioredoxin reductases and peroxiredoxins, to survive within macrophages and cause disease (49–51). Thus, the loss of Rv0148 may destabilize this network, leading to increased oxidative damage and reduced intracellular survival as we reported previously (18).

The redox-sensing transcriptional landscape in *Mtb* includes regulators such as MosR and WhiB family proteins (49, 52). MosR, a homolog of OhrR, senses oxidative stress and derepresses protective genes, while WhiB6 modulates ESX-1 expression and virulence gene transcription in response to redox changes(36, 49, 53). Furthermore, the WhiB proteins contain iron-sulfur clusters that undergo oxidation-reduction reactions, altering their DNA-binding activity and transcriptional output (54, 55). Therefore, in the *Δrv0148* mutant, the altered redox environment may affect the oxidation state of these clusters, leading to dysregulated gene expression. Additionally, expression of PknG, a serine/threonine kinase involved in redox signaling and metabolic regulation was downregulated in the Rv0148 mutant, which may further contribute to the observed proteomic shifts related to stress response and intracellular survival of Mtb (56, 57). Taken together, the interplay between Rv0148 and these redox-sensitive regulators suggests a complex network of transcriptional control that integrates environmental cues to modulate *Mtb* physiology to resist the host antimicrobial response

Our data also indicates that post-translational modifications, particularly acetylation, were altered in the *Δrv0148* mutant. Acetylation of stress-related and ESX-1 proteins suggests a broader reprogramming of the proteome. Protein acetylation has been linked to *Mtb* virulence, drug resistance, and metabolic regulation, with acetyltransferases such as Eis and Pat modulating key survival pathways (58–62). For example, Eis acetylates aminoglycosides and contributes to antibiotic resistance, while Pat regulates central metabolism and stress adaptation in Mtb (61–64). The observed shifts in acetylation targets in the *Δrv0148* mutant may reflect changes in metabolic flux or redox state, further implicating Rv0148 in post-translational control mechanisms. Moreover, acetylation of ribosomal proteins and chaperones can influence protein stability and folding efficiency, potentially exacerbating proteotoxic stress in *Δrv0148* mutant (65, 66). These findings suggest that Rv0148 may indirectly regulate acetylation dynamics through its impact on cellular redox balance.

We observed differential expression of peptidoglycan-associated proteins, which indicate cell wall remodeling in *Δrv0148* mutant. Particularly, upregulation of enzymes such as FhaA and IniA, involved in cell wall repair and remodeling, suggests an adaptive response of *Δrv0148* mutant to maintain envelope integrity under stress (67–69). Conversely, downregulation of biosynthetic components may compromise cell wall robustness, increasing susceptibility of *Δrv0148* mutant to antibiotics and host defenses as we previously reported (18). The peptidoglycan layer of *Mtb* has unique modifications with high levels of cross-linking and amidation that confer resistance to lysozyme and antimicrobial peptides (70, 71). Disruption of these modifications in ΔRv0148 may impair structural integrity and immune evasion. Additionally, cell wall remodeling is tightly linked to *Mtb* dormancy and reactivation, with enzymes such as LdtMt2 and MurA playing roles in latent infection and antibiotic tolerance (72, 73). Thus, altered expression of these enzymes in Δrv0148 mutant suggests that Rv0148 may influence *Mtb*’s transition between replicative and non-replicative states, with implications for bacterial persistence and treatment outcomes.

Finally, our proteomic analysis of *Δrv0148* mutant reveals a complex interplay between redox stress, secretion, translation, and cell wall remodeling in *Mtb*’s adaptation to survive in hostile condition. Based on our data we propose a model in which Rv0148 functions as a central integrator of environmental cues linking those biological functions. Thus, deletion of Rv0148 triggers a compensatory but incomplete stress response, characterized by overproduction of chaperones and select secretion components, alongside suppression of antioxidant enzymes and ribosomal proteins. This imbalance may compromise oxidative stress resistance, secretion fidelity, and cell envelope stability, ultimately attenuating virulence of Mtb.

Nonetheless, our studies have some limitations. First, the proteomic dataset may not capture low-abundance or transient proteins, and our conclusions are based on correlative associations rather than direct mechanistic assays. Second, the redox-sensitive regulatory hypothesis remains speculative without biochemical evidence of Rv0148’s sensing mechanism or interaction partners that contributes to the phenotype of *Δrv0148* mutant. Finally, while we observed changes in acetylation-associated proteins, targeted post-translational modification mapping would be required to confirm these as functionally relevant.

## Data availability

The raw and processed data generated and/or analyzed during the current study are presented in supplementary tables 1 and 2 of this manuscript.

## Author contributions

K.P and G.B initiated the study of proteomics. G.B performed the experiments. AD arranged the Mass spectrometry studies and data curing. S.S. and G.B performed data analysis, interpretation and creation of figures and tables. G.B. wrote the draft manuscript.

K.P and S.S supervised and secured the funding for the study. All the authors have read, reviewed, edited and approved the final manuscript for submission.

## SUPPLEMENTARY INFORMATION.

**Supplementary Figure-1.** Network interaction analysis. (**A).** Interaction map of various biological processes comprised of significantly upregulated proteins in *rv0148* mutant compared to WT. (**B**). Interaction map of various biological processes comprised of significantly downregulated proteins in *rv0148* mutant compared to WT.

**Supplementary Figure-2.** STING analysis of significantly upregulated (**A)** and downregulated (**B**) proteins in *rv0148* mutant compared to WT showing the interaction among proteins of different categories shown in different colors.

**Supplementary Table-1.** Raw protein expression data for WT and *rv0148* mutant strains generated and used in this study.

**Supplementary Table-2.** List of annotated Mtb proteins and their absolute expression levels in WT and *rv0148* mutant. Data in these tables were used to generate all the figures shown in this article.

